# An objective-based prioritization approach to improve trophic complexity through ecological restoration

**DOI:** 10.1101/2021.03.09.434521

**Authors:** Emma Ladouceur, Jennifer McGowan, Patrick Huber, Hugh Possingham, Davide Scridel, Roel van Klink, Peter Poschlod, J. Hans C. Cornelissen, Costantino Bonomi, Borja Jiménez-Alfaro

## Abstract

1. Reassembling ecological communities and rebuilding habitats through active restoration treatments requires curating the selection of plant species to use in seeding and planting mixes. Ideally, these mixes should be assembled based on attributes that support ecosystem function and services, promote plant and animal species interactions and ecological networks in restoration while balancing project constraints. Despite these critical considerations, it is common for species mixes to be selected opportunistically. Reframing the selection of seed mixes for restoration around ecological objectives is essential for success but accessible methods and tools are needed to support this effort.
2. We developed a framework to optimize species seed mixes based on prioritizing plant species attributes to best support different objectives for ecosystem functions, services, and trophic relationships such as pollination, seed dispersal, and herbivory. We compared results to approaches where plant species are selected to represent plant taxonomic richness, dominant species, and at random. We tested our framework for 176 plant species found in European alpine grasslands and identified 163 associated attributes affiliated to trophic relationships, ecosystem functions, and services.
3. In all cases, trophic relationships, ecosystem functions, and services can be captured more efficiently through objective-based prioritization using the functional identity of plant species. Solutions (plant species lists) can be compared quantitatively, in terms of costs, species, or objectives. We confirm that a random draw of plant species from the regional plant species pool cannot be assumed to support other trophic groups and ecosystem functions and services.
4. *Synthesis and Applications*. Our framework is presented as a proof of concept to help restoration practitioners better apply quantitative decision–support to plant species selection in order to meet ecological restoration outcomes. Our approach may be tailored to any restoration initiative and habitat where seeding or planting mixes will be applied in active treatments. As global priority and resources are increasingly placed into restoration, this approach could be advanced to help make efficient decisions for many stages of the restoration process.

## Introduction

The specific objectives of terrestrial ecological restoration will vary, but generally aim to return a habitat to a naturally functioning and stable state. Restoration is often operationalized through the planting or seeding of mixtures of plant species as an active treatment meant to re-establish plant communities in degraded sites, usually informed by the plant species composition at reference sites (Brudvig & Mabry, 2008; Zobel et al., 1998). This begins by defining a species pool for the ecosystem of interest (the regional species pool, Zobel et al., 1998), and by taking stock of what species can be sourced from the wild or from commercial native seed producers (the restoration species pool) (Ladouceur et al., 2018; Zobel et al., 1998). However, seed and planting mixes used for restoration are often a low diversity subset of the relevant species pool, and are composed opportunistically (Barr et al., 2016). Species selection must be balanced within project constraints (eg. budgets, labour, time), and within other project targets (eg. increase plant cover, prevent erosion). These species mixes have a major impact on restoration success and have an impact on the multi-taxa functionality of the restored ecosystems (Guiden et al., 2021). How species mixes can be optimized to maximise restoration goals efficiently within project constraints remains an open question and an urgent task for implementing the United Nations Decade on Ecosystem Restoration.

It is widely recognized that rebuilding habitats requires the consideration of ecosystem services and functions, fauna, and plant-animal relationships (Kollmann et al., 2016; McAlpine et al., 2016). Integrating these relationships in ecological restoration is a complex task that remains largely unaddressed despite increased calls for consideration (Cross et al., 2020; Dixon, 2009; Lindell, 2008; Majer, 1989, 2009; Menz et al., 2011). Plant functional traits can help identify ecosystem services or functions facilitated by plant species, and can lead to favourable restoration outcomes (Brudvig & Mabry, 2008; Zirbel et al., 2017). Fauna also contribute crucial ecosystem functions to plants, such as seed dispersal (regeneration), pollination (seed production), herbivory (reduction of competitive dynamics), and patchy nutrient return (Olff & Ritchie, 1998). Optimizing plant species mixes to facilitate multiple ecosystem services and functions, including those performed by other trophic levels, could thus enhance restoration success, as shown by the establishment of fruit bearing trees to facilitate dispersal from other diverse patches by frugivores in tropical rainforests (Heelemann et al., 2012; Lamb et al., 1997).

However, the restoration of ecosystem services, functions, and animal communities is challenging due to complex processes, life cycles, and dependence on plants as well as other trophic levels (Chan et al., 2006; Guiden et al., 2021). Plant-animal interaction networks, both mutualistic (pollination and frugivory) and antagonistic (herbivory) are highly non-random (Bascompte & Jordano, 2007; Lewinsohn et al., 2006; Rezende et al., 2007), and a disruption in these interactions can lead to trophic cascades across and within systems (Knight et al., 2005; Valiente-Banuet et al., 2015). Plant-animal interaction networks are often nested, that is, some species have many interactions in their networks, and many species have few (Bascompte et al., 2003). When considering balancing project constraints and restoration targets in a relatively low-diversity species mix for restoration, it is unlikely that a random draw from the regional plant species pool will provide resources to optimize trophic networks and other ecosystems services and functions. Systematic decision-making can quantitatively support complex multivariate decision-making problems such as this (Chan et al., 2006; Hill et al., 2014; M’Gonigle et al., 2016).

Here, we present a proof-of concept for the optimization of active restoration species mixtures (for seeding or planting treatments) for supporting different objectives. We used species-rich European subalpine and alpine calcareous grasslands as a case study. These habitats are sensitive to disturbance, and impacted by ski resorts and other tourism activities, making them a target system for ecological restoration across European Natura 2000 sites (Garcia-Gonzalez, 2008, p.). We identified 176 plant species that frequently occur in the target ecosystem on a biogeographical scale as the potential regional and restoration species pool of interest (Ladouceur et al., 2018; Zobel et al., 1998). We used trait databases and literature to compile traits related to regeneration and relationships between the 176 plant species in our species pool and the insects, birds and mammals that are typical of these habitats and depend on particular plant species for various life stages. Hereafter, we refer to the traits and aspects of plant species that represent these relationships and characteristics of interest, as plant attributes.

Our primary aim is to develop and evaluate a quantitative decision-making framework to assist in species selection for seeding and planting mixes for restoration projects. To do so, we designed five objectives for prioritizing plant attributes that support ecosystem functions, services and trophic dependencies. We optimized for these objectives by finding the smallest number of plant species needed to deliver all of the attribute targets set within each objective (Possingham et al., 2000). We then developed four plausible species selections to compare with prioritized selections including a focus on dominant species, random draws from the plant species pool to represent different taxonomic resolutions, and completely random draws from the plant species pool. We used our study system to investigate whether optimized species pools deliver objectives for ecosystem functions or services more efficiently than selecting dominant plant species, for taxonomic richness, or randomly.

## Methods

We designed and tested an optimization approach to prioritize species mixes for planting or seeding in restoration projects based on ecological objectives. Below, we describe how we 1) selected plant species from a defined regional species pool; 2) identified plant attributes; 3) constructed objectives and optimized attributes; and 4) evaluated across approaches.

### Species selection

We compiled a list of the most frequent native species occurring in alpine calcareous grassland habitat types on a continental scale, using a synthesis of >1 million field surveys (Schaminée et al., 2016), reporting species frequencies in the habitat types of the European habitat classification system (EUNIS, www.eunis.org), directly assigned to habitat types of conservation concern (Table S1). We identified native plant species that occur above a particular frequency (>5% of total occurrences) in calcareous alpine grassland habitat types on a European-wide scale. Expert opinion suggests that species below this frequency were found to be more typical of other habitat types. This resulted in a list of 176 native plant species that occur frequently in the calcareous alpine grasslands of continental Europe. We considered this to be the species pool of this habitat and we assumed all species can co-occur or can co-exist. Further, we consider all species in the pool as equal candidates for inclusion in seed mixes to meet prioritization objective targets.

### Attribute selection

For the 176 plant species that were of interest for our goals, we collated traits related to dispersal, phenology, and nitrogen fixation available from the TRY plant trait database (Kattge et al., 2011), as well as associations with mammals, birds, and herbivorous and pollinating insects from additional sources (see Table 1). The list of associated faunal species was refined to keep only species that occur in this habitat. Plant species frequency of occurrence values were used to rank plant species’ relative abundance within the habitat type on a biogeographic scale, which we used to classify plant species dominance for a fixed species list for comparison with prioritized objectives (Table 1, Table S1).

**Table 1:**
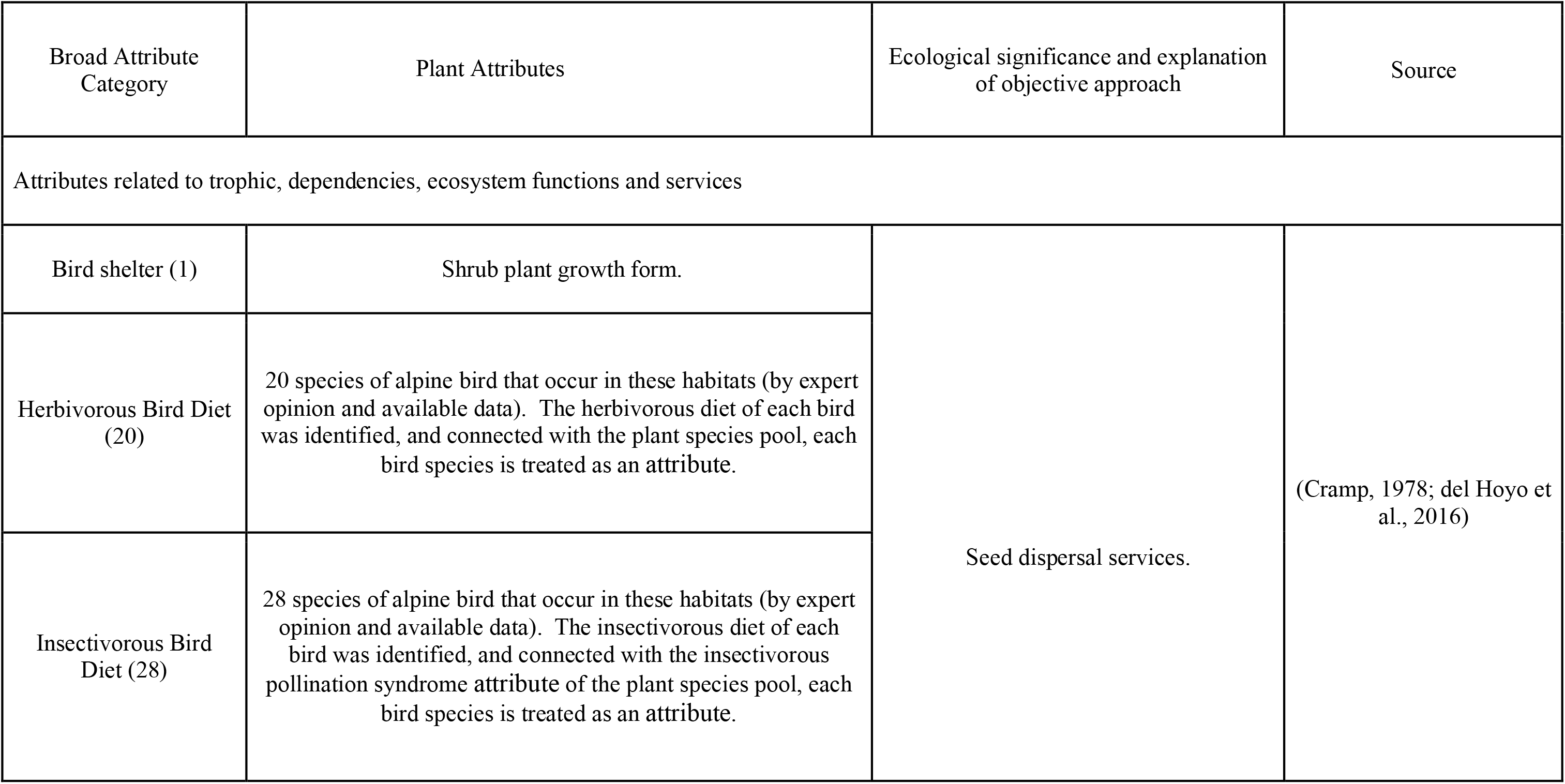
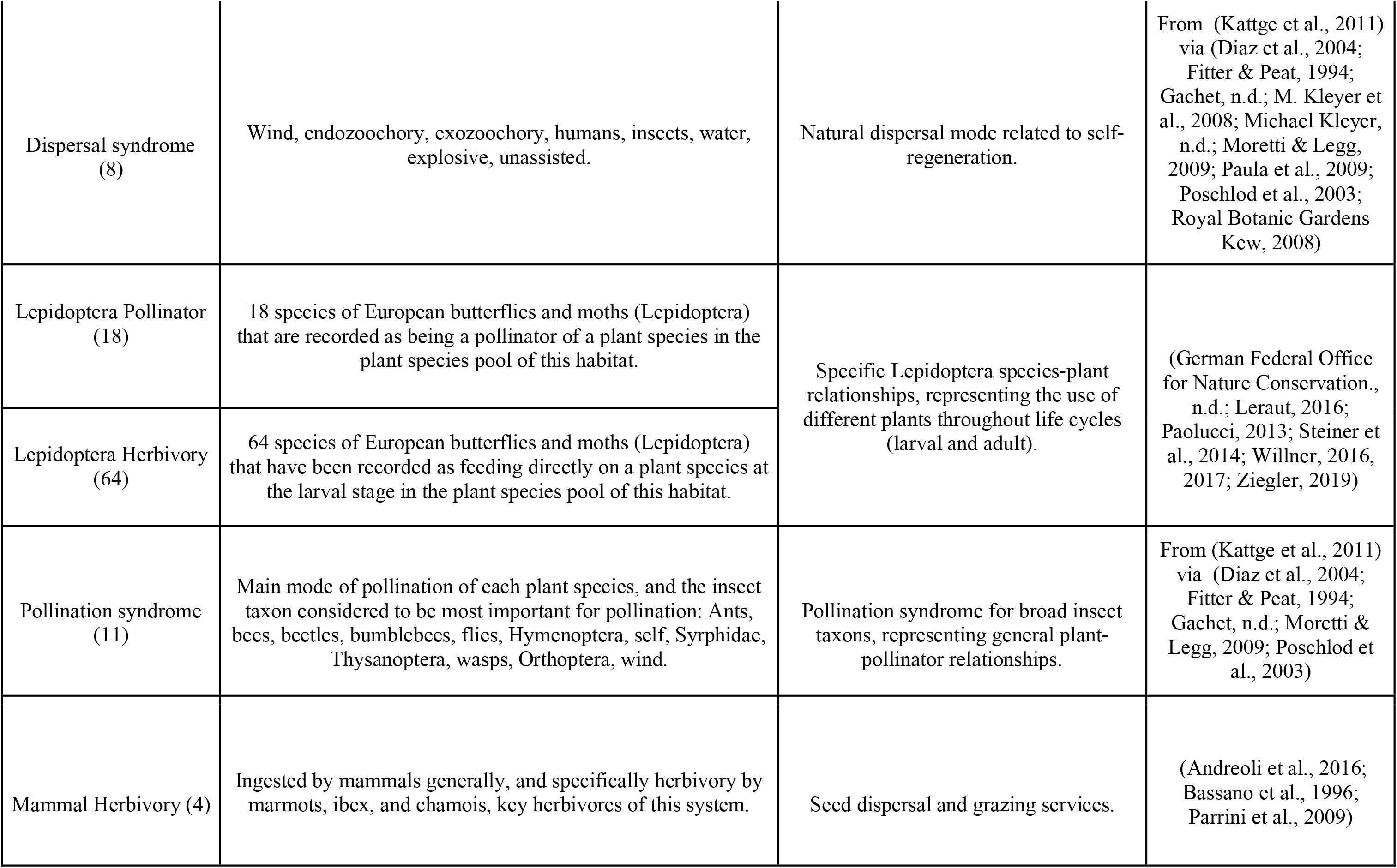
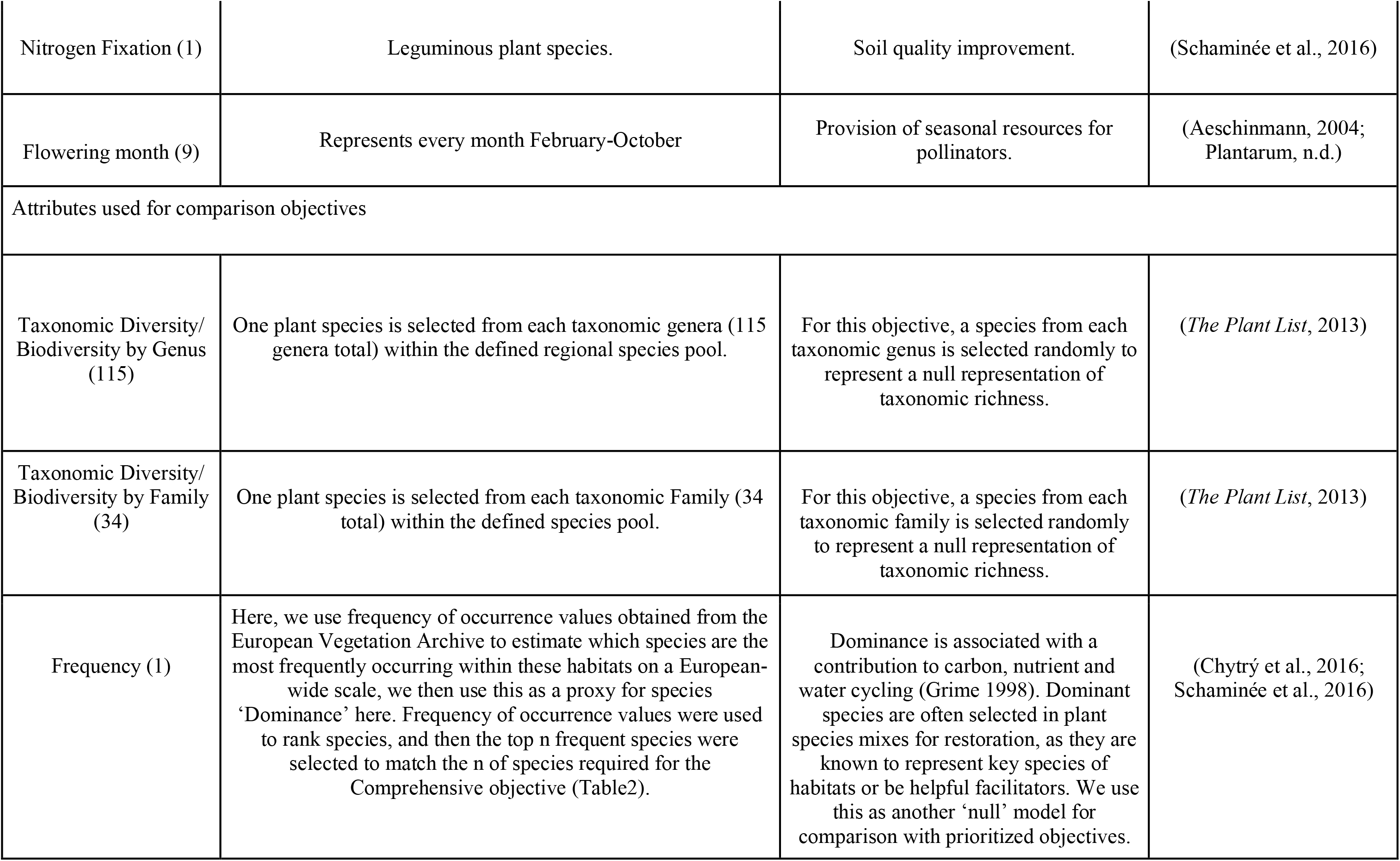
Plant species attributes, ecological role and data source of each attribute used, grouped by broad ecosystem function or service category. Complete list in Table S1.

| Broad Attribute Category | Plant Attributes | Ecological significance and explanation of objective approach | Source |
| --- | --- | --- | --- |
| Attributes related to trophic, dependencies, ecosystem functions and services |  |  |  |
| Bird shelter (1) | Shrub plant growth form. | Seed dispersal services. | (Cramp, 1978; del Hoyo et al., 2016) |
| Herbivorous Bird Diet (20) | 20 species of alpine bird that occur in these habitats (by expert opinion and available data). The herbivorous diet of each bird was identified, and connected with the plant species pool, each bird species is treated as an attribute. |  |  |
| Insectivorous Bird Diet (28) | 28 species of alpine bird that occur in these habitats (by expert opinion and available data). The insectivorous diet of each bird was identified, and connected with the insectivorous pollination syndrome attribute of the plant species pool, each bird species is treated as an attribute. |  |  |
| Dispersal syndrome<br>(8) | Wind, endozoochory, exozoochory, humans, insects, water, explosive, unassisted. | Natural dispersal mode related to self-regeneration. | From (Kattge et al., 2011) via (Diaz et al., 2004; Fitter & Peat, 1994; Gachet, n.d.; M. Kleyer et al., 2008; Michael Kleyer, n.d.; Moretti & Legg, 2009; Paula et al., 2009; Poschlod et al., 2003; Royal Botanic Gardens Kew, 2008) |
| Lepidoptera Pollinator<br>(18) | 18 species of European butterflies and moths (Lepidoptera) that are recorded as being a pollinator of a plant species in the plant species pool of this habitat. | Specific Lepidoptera species-plant relationships, representing the use of different plants throughout life cycles (larval and adult). | (German Federal Office for Nature Conservation., n.d.; Leraut, 2016; Paolucci, 2013; Steiner et al., 2014; Willner, 2016, 2017; Ziegler, 2019) |
| Lepidoptera Herbivory<br>(64) | 64 species of European butterflies and moths (Lepidoptera) that have been recorded as feeding directly on a plant species at the larval stage in the plant species pool of this habitat. |  |  |
| Pollination syndrome<br>(11) | Main mode of pollination of each plant species, and the insect taxon considered to be most important for pollination: Ants, bees, beetles, bumblebees, flies, Hymenoptera, self, Syrphidae, Thysanoptera, wasps, Orthoptera, wind. | Pollination syndrome for broad insect taxons, representing general plant-pollinator relationships. | From (Kattge et al., 2011) via (Diaz et al., 2004; Fitter & Peat, 1994; Gachet, n.d.; Moretti & Legg, 2009; Poschlod et al., 2003) |
| Mammal Herbivory (4) | Ingested by mammals generally, and specifically herbivory by marmots, ibex, and chamois, key herbivores of this system. | Seed dispersal and grazing services. | (Andreoli et al., 2016; Bassano et al., 1996; Parrini et al., 2009) |
| Nitrogen Fixation (1) | Leguminous plant species. | Soil quality improvement. | (Schaminée et al., 2016) |
| Flowering month (9) | Represents every month February-October | Provision of seasonal resources for pollinators. | (Aeschinmann, 2004; Plantarum, n.d.) |
| Attributes used for comparison objectives |  |  |  |
| Taxonomic Diversity/<br>Biodiversity by Genus<br>(115) | One plant species is selected from each taxonomic genera (115 genera total) within the defined regional species pool. | For this objective, a species from each taxonomic genus is selected randomly to represent a null representation of taxonomic richness. | (*The Plant List*, 2013) |
| Taxonomic Diversity/<br>Biodiversity by Family<br>(34) | One plant species is selected from each taxonomic Family (34 total) within the defined species pool. | For this objective, a species from each taxonomic family is selected randomly to represent a null representation of taxonomic richness. | (*The Plant List*, 2013) |
| Frequency (1) | Here, we use frequency of occurrence values obtained from the European Vegetation Archive to estimate which species are the most frequently occurring within these habitats on a European-wide scale, we then use this as a proxy for species 'Dominance' here. Frequency of occurrence values were used to rank species, and then the top n frequent species were selected to match the n of species required for the Comprehensive objective (Table2). | Dominance is associated with a contribution to carbon, nutrient and water cycling (Grime 1998). Dominant species are often selected in plant species mixes for restoration, as they are known to represent key species of habitats or be helpful facilitators. We use this as another 'null' model for comparison with prioritized objectives. | (Chytrý et al., 2016; Schaminée et al., 2016) |

We then grouped the 163 plant attributes into nine broad categories based on the ability to support specific ecosystem functions or services (Table 1): bird trophic diet, bird herbivory, bird shelter, seed dispersal syndrome, Lepidoptera relationships (species specific-pollination, herbivory), pollination syndrome, mammal herbivory, nitrogen fixation, and flowering month. The range of attributes supported by plant species varied greatly, with some highly specialized plant species supporting only one attribute (e.g., *Galium estebanii*) while others support many attributes (e.g., *Poa alpina,* alpine meadow grass (56 attributes) and *Sedum album,* white stonecrop (58 attributes)) (Table S1).

To assign attributes to species we used a binary classification scheme, where a value of 1 was used when an attribute was present in a given plant species. In some instances, the presence of an attribute is dependent on the connection between the plant species and a species of other trophic groups, such as birds or butterflies (Lepidoptera). Some Lepidoptera species depend on different plant species at different life stages (larval herbivory vs adult pollination/visitation), which we accounted for (Table 1). For birds, we connected trophic dependencies between attributes. For example, the plant species *Aster alpinus* (alpine aster) has a beetle pollination syndrome, and the bird *Turdus torquatus* (ring ouzel) feeds on beetles as part of its diet, so alpine aster is potentially an important habitat component for beetles, and the ring ouzel (see Table S1).

### Objective Construction, Comparison lists and Prioritization

We constructed five objectives for prioritizing species based on setting targets which deliver: 1) representation of all attributes present in the species pool (“Comprehensive” (N=163 attributes)); 2) specific processes and taxa that play key roles in ecosystem regeneration, specifically, species-specific seed dispersal and pollination for birds (“Bird”, N= 48 attributes) and 3) Lepidoptera Relationships (pollination and herbivory) (“Lepidoptera Relationship”, N= 82 attributes), 4) representation of both levels of taxonomic plant richness in combination with Lepidoptera relationships (“Pairwise Lepidoptera + Plant Rich Family”, N=116 attributes including plant families counted as an attribute), 5) (“Pairwise Lepidoptera + Plant Rich Genus”, N= 197 attributes). We compared these five objectives to four comparison lists - plant species selections meant to serve as plausible opportunistic approaches for creating species mixes. These include; 1) a fixed list of the most frequent species occurring in these habitats at a biogeographic scale, as a proxy for dominant species (“Dominants”, N= a fixed list of 37 plant species), 2) a representation of plant diversity through taxonomic richness at the 2) family (“Plant Rich Family”, N= 34 families), and 3) genus level (“Plant Rich Genus”, N= 115 genera); and 4) selecting plant species at random (“Random”) (see Table 2). To compare a species-mix of dominant species to prioritized objectives, we sorted dominant species by frequency of occurrence values, and created a fixed list of ‘dominant’ species equal to the number of species in the Comprehensive solutions for direct comparison between the performance of the two species lists in terms of representing attributes that potentially support particular ecosystem functions and services. We consider a single presence to be sufficient to capture the attributes.

**Table 2:** Summary of how the nine ecological restoration prioritization objectives were implemented in the decision support software prioritizr, showing the details of the attributes within each objective, and the number of attributes targeted to be included in the selection of plant species for each objective. Each prioritized objective (Comprehensive, Bird, Lepidoptera Relationship, Pairwise Lepidoptera + Plant Rich Family, Pairwise Lepidoptera + Plant Rich Genus) aims to select the fewest number of plant species, while meeting all objective targets. These five prioritized objectives are then compared to four species selections that can serve as a null comparison (Dominants, Plant Rich Family, Plant Rich Genus, Random).

| Objective | Attributes Description | Objective targets |
| --- | --- | --- |
| Prioritized Objectives |  |  |
| Comprehensive | 163 attributes, including all attributes from Table 1 specified to the species level. | Ensure all 163 single species-specific attributes are represented at least once |
| Bird | 49 attributes representing 28 species of alpine grassland birds, under three broad ecosystem function categories: herbivores (20 bird sp.), insectivores (28 bird sp.), and bird shelter (1 attribute) | Ensure all 49 bird related attributes are represented at least once |
| Lepidoptera Relationship | 82 attributes representing 76 unique species of butterflies and moths (Lepidoptera) under two broad ecosystem function categories: pollinators (18), larval (64). 6 Lepidoptera species have multiple life-stage requirements represented in the dataset. | Ensure all 82 species-specific Lepidoptera relationship related attributes are represented at least once |
| Pairwise Lepidoptera +<br>plant rich Family | 34 plant taxonomic families + 82 plant-pollinator relationships as<br>attributes | Include 1 plant species belonging to every family,<br>and all 49 plant-pollinator related attributes are<br>represented at least once |
| Pairwise Lepidoptera +<br>plant rich Genus | 115 plant taxonomic genera + 82 plant-pollinator relationships as<br>attributes | Include 1 plant species belonging to every single<br>genus, and all 82 plant-pollinator related<br>attributes are represented at least once |
| Comparison Objectives |  |  |
| Dominants | Species frequency of occurrence on a biogeographical scale identified<br>and rank ordered in terms of dominance (Chytrý et al. 2016; Schaminée<br>et al. 2016). | Include the n most frequent plant species to match<br>n species required for the Comprehensive<br>selection, as a single fixed list to allow direct<br>comparison. |
| Plant Rich Family | 34 plant taxonomic families in the dataset. | Include 1 randomly selected plant species<br>belonging to every family to represent 'plant<br>biodiversity' at the Family level. |
| Plant Rich Genus | 115 plant taxonomic genera in the dataset | Include 1 randomly plant species belonging to<br>every genus to represent 'plant biodiversity' at the<br>genus level. |
| Random | Select plant species randomly in intervals of 5, ranging from 5-120<br>species . | No attribute targets. |

To efficiently find the smallest number of plant species that met the target based-objectives (Table 1), we used the ‘minimum-set’ problem formulation which is commonly applied to spatially-explicit decision making that cost-efficiently meet targets for conservation features (e.g. habitats, species ranges, or ecological processes) (Possingham et al., 2000). We adapted inputs to apply it to our non-spatial problem (Hill et al., 2014) (see Supporting Information Appendix 1). To do so, we replaced geographic spatial units with individual plant species, and replaced the features found in those geographic units with the functional attributes assigned to each species, resulting in a plant species-attribute matrix (Figure 1). Each plant species had a unique set of attributes, each attribute with a binary value of ‘0’ or ‘1’, and these values were summed to produce a ‘attribute sum’ for every plant species; that is - the number of attributes that characterise each plant species. For each objective, complementary sets of plant species were identified where collective attributes achieved the minimum targets set (where we considered a minimum target of 1) (Table 2). We set equivalent costs across species (value of 1) so that we could test the outcomes of prioritizing plant species across different objectives independent of costs.

**Figure 1:**
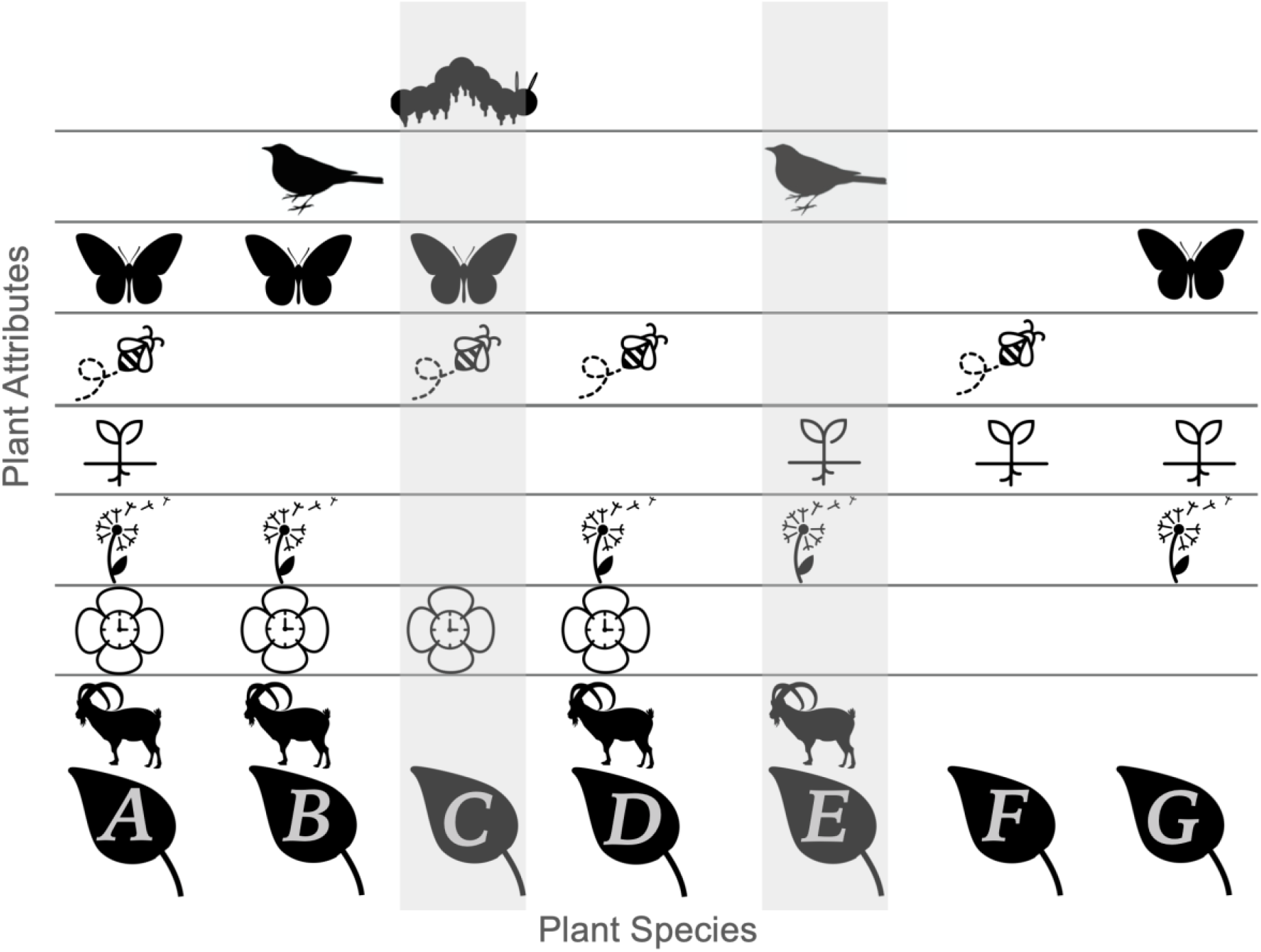
Conceptual framework of plant species prioritization for attributes that enable ecosystem services: a simplified example. Every plant species A-G represented in each vertical column has a unique combination of attributes in the plant species-attribute matrix which are represented in each horizontal row. Plant species C & E capture each attribute just one time when selected together. All images sourced from The Noun Project.

Once objectives were set, all problems were solved using the R package prioritizr (Hanson et al., 2019), with Gurobi 9.0 as an algorithmic solver (Gurobi Optimization Inc., 2018). For each prioritized objective, we set problems in prioritizr with an optimality gap of zero, and the ‘add_gap_portfolio’ function to produce a portfolio of 100 different solutions, where the first solution is the optimal solution to the original data formulation, and every solution thereafter meets targets within the pre-specified optimality gap. This relative gap specifies a threshold worst-case performance for solutions in the portfolio, so in this case, we chose to accept 100 solutions no matter the performance relative to the optimal (gap=0). For all random solutions for comparison, we used ‘add_shuffle_portfolio’ (instead of the gap portfolio). This randomly reordered data prior to solving problems, so plant species were selected under different data formulations to produce a random selection process.

### Evaluation

In comparing and evaluating approaches to plant species selections for species mixes, prioritizr produces two important outputs: optimal solutions to meet targets for objectives (in this case, a plant species list); the feature representation indicating the number of plant attributes represented by a solution, relatively (to possible maximums) or absolutely (total number). We used the first optimal solution of each objective to compare the attribute sum of the plant species selected (Table S1). We compared the mean values of the attribute sum using a Kruskal-Wallis chi-square test to assess differences in the total number of attributes (attribute sum).

We calculated the selection frequency of plant species across all 100 solutions generated to identify the relative irreplaceability of each plant species within a species mix to meet targets for each objective. Where a plant species had a selection frequency of 100 across solutions, we categorised it as *‘irreplaceable’.* Irreplaceability can be interpreted as an index of the likely overall value of a feature, or in this case a plant species, in achieving an objective (Smith et al., 2018). Where a species was chosen between 1 and 99 times, we categorized it as *‘variable’,* and where it was chosen zero (0) times it was categorized as *‘redundant’.*

We evaluated each objective’s ability to capture the nine broad ecosystem functions and service categories defined in Table 1. To do so, we took the species identified in all 100 solutions for each objective (see supporting information), identified the full list of attributes present (Table S1) and calculated the percentage of attributes captured compared to the total number of attributes possible for selection in each broad attribute category (Table 1).

We compared our results to an ad-hoc selection of species that we approximated using a random species selection process. We selected plant species at random in intervals of 5 (ranging from 5 to 125 plant species) and calculated the proportions of attribute provision captured across the same nine broad ecosystem function categories. We considered the mean (50%), upper (75%) and lower quantiles (25%) of each random species selection across all solutions for comparison across objectives.

All prioritizations, figures and analyses were conducted using the R Studio version 1.3.1056 and R version 4.0 environment and language for statistical computing and graphics (R Core Development Team, 2019).

## Results

Across the five ecological objectives, the number of plant species needed to meet each objective’s targets varied widely. For example, the targets for the “Bird” objective were met with only five plant species, targets for the “Pairwise Lepidoptera + Plant Rich Genus” objective required 119 plant species (Figure 2). We also found high variability in the number of attributes (the attribute sum) captured by the individual plant species selected in the solutions (max:59; min:2) (Table S1). The plant species selected in the Comprehensive” and the “Lepidoptera Relationship” objective had a significantly higher attribute sum overall (Kruskal-Wallis s^2^= 146.68, P= <0.001, Table S3, Figure 2, Figure S1) than found in the other objectives.

**Figure 2:**
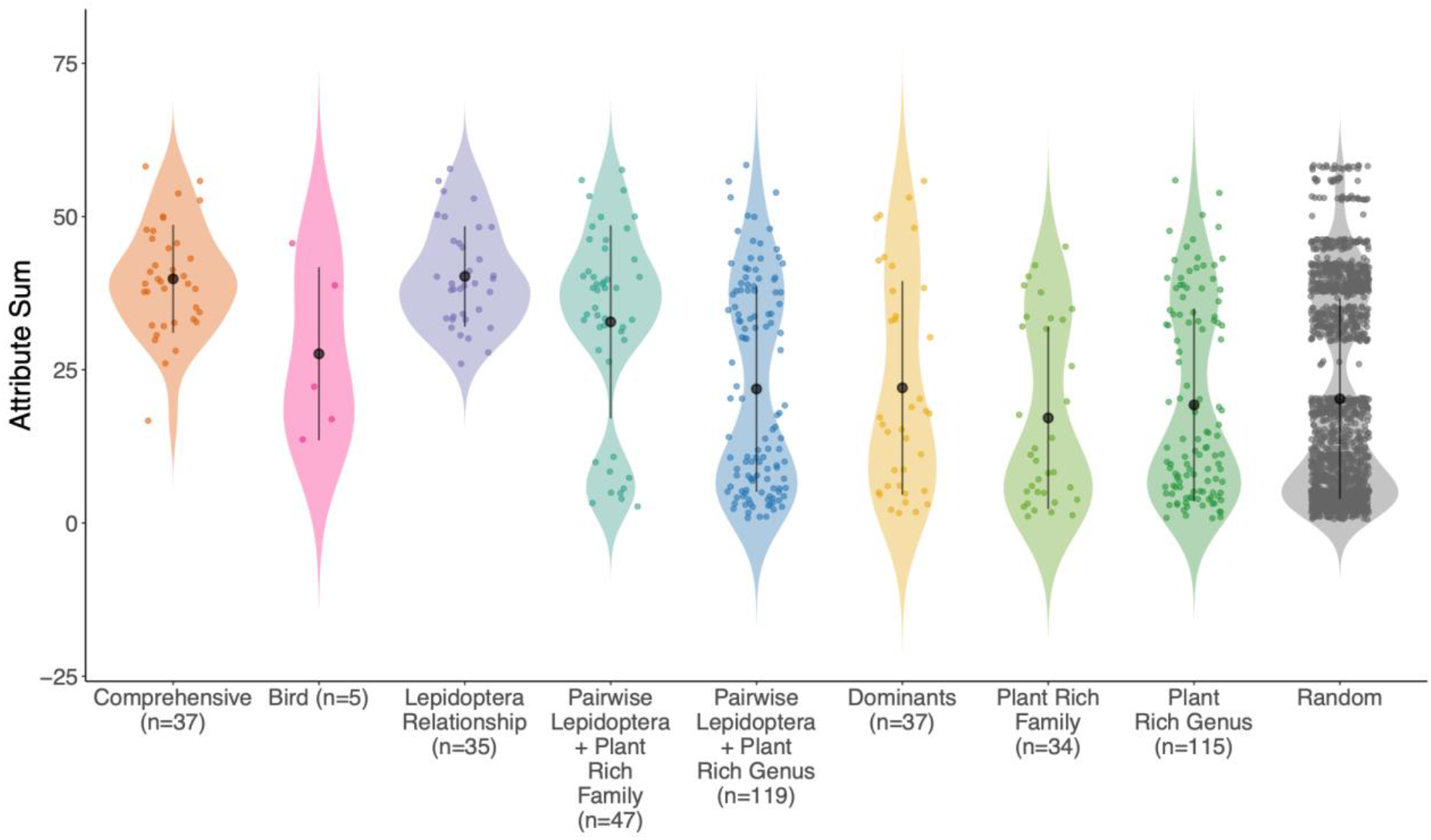
The attribute sum (total number of attributes per plant species) captured by each plant species in the first optimal solution of every objective. Each objective solution is labelled along the x-axis, and reflects the selection of species needed to meet the targets of that objective. Each point is a plant species. The large black points represent the mean and the bars represent the standard error around the mean. The shaded area represents the spread and the density of the data. n: number of plant species in the solution of every objective.

This prioritization approach favours plant species with a high attribute sum, yet also prioritizes plant species that supports unique or rare attributes, as these species may be considered irreplaceable (Figure 3). In the case of the Comprehensive and the Lepidoptera Relationship objective, many plant species were irreplaceable. I contrast, in the bird objective, many plant species were of variable importance, and thus could be interchangeable (Figure 3, Table S4).

**Figure 3:**
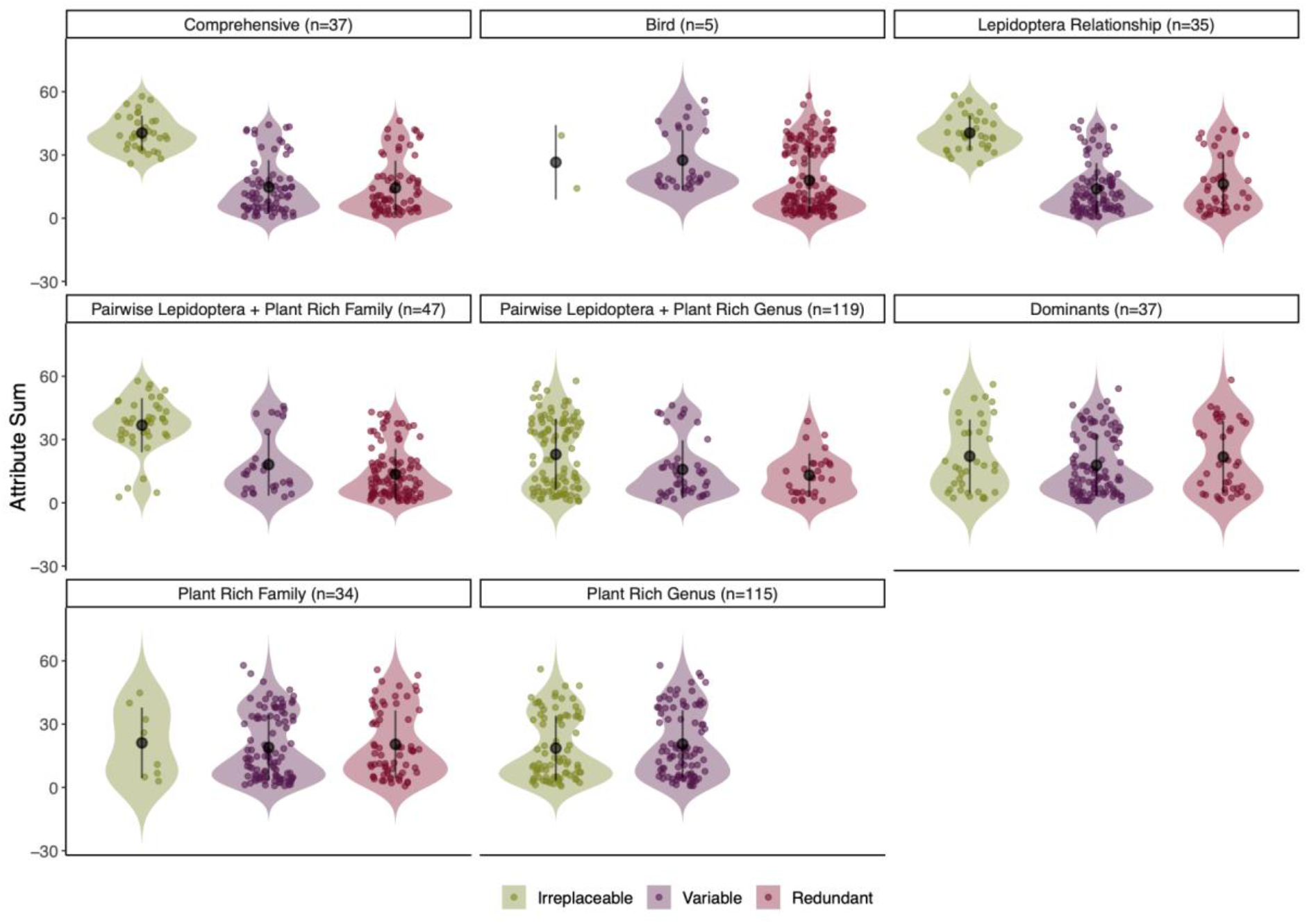
The attribute sum (total number of attributes per plant species) captured by each plant species for each objective, for plants species that were Irreplaceable (Selection Frequency (SF) =100/100), Variable (SF=99>1/100), or Redundant (SF=0). Each point is a plant species. The black points represent the mean and the bars represent the standard error around the mean. The shaded area represents the spread and the density of the data.

Figure 4 illustrates the number of plant species selected across the first solutions for each objective and the percentage of the attributes captured relative to the total number of attributes in each ecosystem function category. This demonstrates the trade-off between the number of plant species selected and the provision of minimum sets of attributes. It also examines how well a single objective captures broad ecosystem function and services compared to the performance of other objectives.

**Figure 4:**
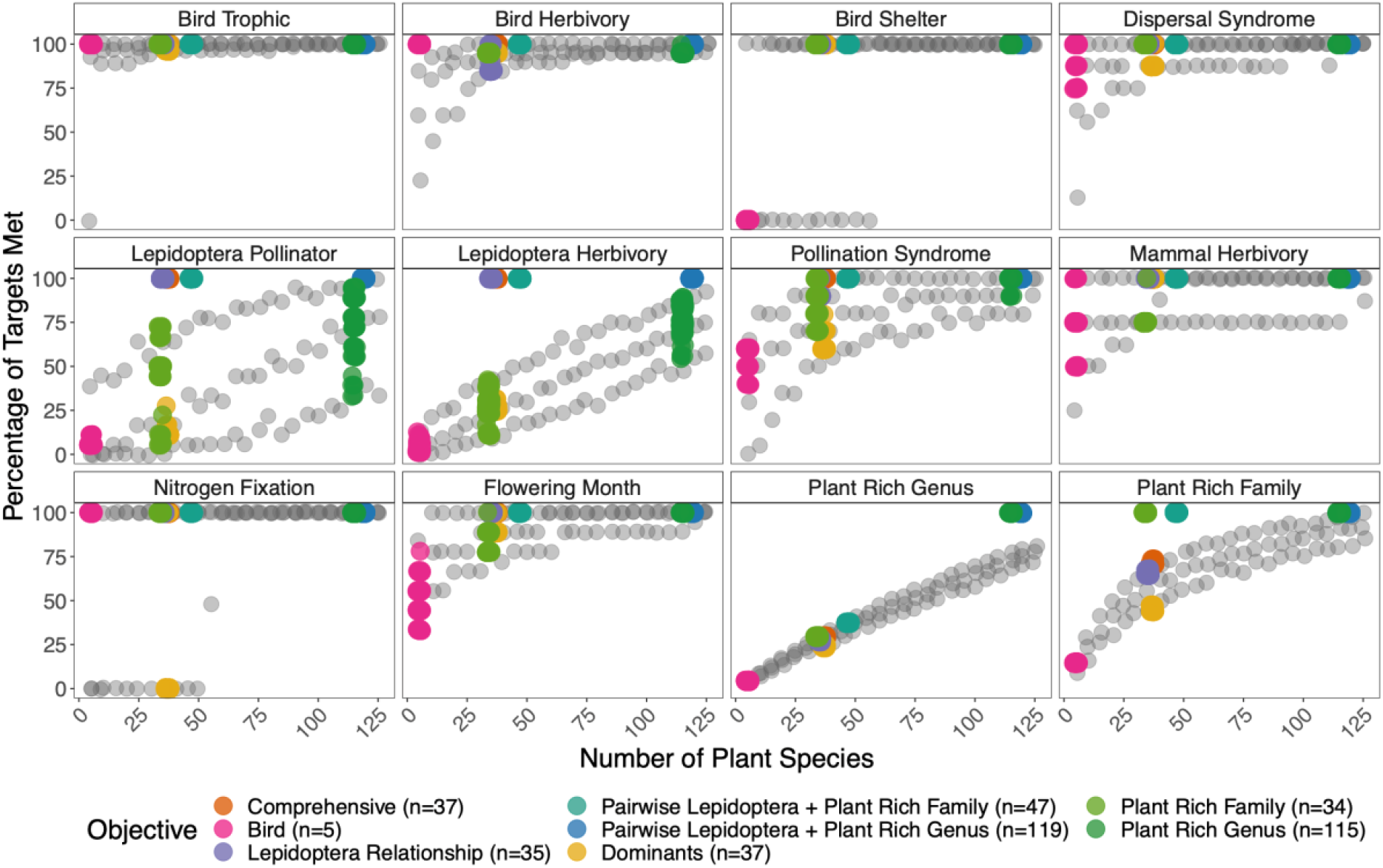
Trade-offs between number of plant species selected for each objective, and the corresponding proportion of ecosystem function targets captured compared across objectives. Optimizations were run 100 times for each targeted coloured objective and all unique runs are shown here as points. Grey points represent the mean, upper (75%) and lower (25%) quantiles of runs for randomly selected species in intervals of 5 from 5-120 plant species.

Objectives that did not set out to prioritize a specific category of ecosystem function had variable performance. For example, the “Bird” objective performed poorly for all plant-pollinator related ecosystem functions, capturing <25% of the attributes needed to support plant-pollinator, nectar, and larval functions (Figure 4). This is unsurprising given the “Bird” objective only needed five plant species to achieve the targets. Alternatively, the “Comprehensive” objective, which aimed to represent each of the 163 attributes found across the entire species pool once (and did so with 37 plant species), met 100% of targets set (1 of every attribute). However, the species prioritized in this objective performed no better than the random species selection for representing Genus and Family levels of plant diversity representation (Fig 4).

Overall, the random selection of plant species performed well for ecosystem functions that are supported by common attributes across plant species (e.g., bird trophic, bird herb, bird shelter), but worse than our prioritization when the ecosystem function is supported by a highly specialized attribute (e.g., plant-pollinator relationships) (Figure S2). In general, the smaller the number of randomly selected plant species, the worse the performance for providing ecosystem functions and services. Even when large numbers of randomly selected plant species are considered, provision of some trophic relationships, or ecosystem function and service groups were found to be low (Fig. S2).

## Discussion

Plant species have a unique combination of functional attributes that contribute to important ecosystem processes and trophic relationships in different ways. Here, for the first time, we have developed and tested an approach for prioritizing plant species in order to represent multiple plant attributes that potentially support trophic complexity and ecosystem services and functions in species mixes for active restoration treatments. Our results show that species selection approaches targeting for taxonomic richness, dominant species and/or with a random approach may not support higher trophic levels and the ecosystem functions and services they provide as efficiently as our objective-based approaches. Critically, our results illustrate that higher trophic levels and ecosystem functions can, in some instances, be supported well when plant species richness is relatively low. Conversely, trophic relationships and ecosystem functions can in some cases be unsupported and low while plant species richness is high. We confirm that a random draw of plant species from the regional plant species pool cannot be assumed to support other trophic groups and ecosystem functions and services. This has important implications for the design and implementation of species mixes for restoration projects which aim to reach multiple restoration objectives such as plant diversity, higher trophic levels and certain ecosystem functions and services tied to plant species identities.

### Prioritizing functional attributes

Some ecosystem functions and services are captured by plant species selections easily, even when these are not the targets of the objective. In these cases, the functional plant attribute is abundant (eg. wind dispersal syndrome) within the plant species pool. For example, bird diets are often generalised to a plant genus or family (eg. Asteraceae), so minimum diet requirements for the bird species represented here do not require many plant species to meet minimum provisional targets. By randomly selecting species from the species pool, these attributes are often captured in a minimum amount of plant species. The bird objective only represents five plant species to provide a minimum diet for twenty-eight species of alpine birds, and in practice a species mix designed for birds would benefit from higher representation of these plant functional attributes and diet options.

In other cases, where a specialist relationship between an attribute and a plant species exists, targets are not captured well, unless an objective is prioritized for such. For example, relationships between plants and insect herbivores are often specialised, making many plant species irreplaceable when optimizing the plant community for herbivores. The objectives for plant taxonomic richness, for dominant species and for randomly selecting species do not meet minimum targets for plant-pollinator relationships, even when up to 125 plant species are selected. The fewer plant species that are selected, the higher the risk that resources for herbivores and pollinators will not be provided within the plant species mix. However, when targeted, all Lepidoptera species relationships with particular plant species in terms of larval herbivory or pollination (82 Lepidoptera relationships total) can be represented at least once within a species mix with 35 targeted plant species. Negative changes within ecosystems can lead to trophic cascades (Knight et al., 2005), and in restoration, there is the opportunity to directly support these connections between organisms positively facilitate regeneration processes and ecological networks (Harvey et al., 2017; Valiente-Banuet et al., 2015) through this framework. When considered this way, one can ask if the species pool used in restoration is providing adequately for the species pool of other trophic levels within that habitat while balancing multiple targeted outcomes.

Conversely, depending on how plant taxonomic diversity is defined (representing one species from every taxonomic Family or Genus), it is not always represented well by objectives prioritized for attributes, or by randomly selecting species, but can be captured efficiently through targeted selection. Additionally, both attributes and plant biodiversity can be captured efficiently together when both are set as targets (Pairwise objectives). Seed mixes matter for restoration success and can be optimized according to many factors (Barr et al., 2016), but require the balancing of multiple targets which is a complex multivariate decision-making task that can make use of decision-support tools as demonstrated here.

Additionally, seeding and planting treatments for restoration are restricted by many confounding constraints including budgets, labour, and project size and so restoration species mix treatments are often quite low (Barr et al., 2016). Where constraints are present, prioritizing plant species to optimise particular targets can be a potentially beneficial method to decide which species to include in low diversity treatments. This method has similarities to methods for filtering plant species lists based on particular targets (Brudvig & Mabry, 2008), but offers the unique advantage of optimising targets according to constraints, and offers quantitative support for comparing different options easily both in terms of targets and cost.

### Indications and Further Development

In order to test this proof of concept, it was necessary to make some simplifying assumptions. Focusing on a study system with relatively good knowledge on frequently occurring plant species, we selected attributes based on available data in the target system. Rare plant species, which have not been thoroughly studied, are often documented as having few attributes, resulting selection bias towards representing common species. A prioritized solution can only be as good as the data available, and the prioritization objectives set out here are limited by the available data. Generalised data on pollination syndromes or seed dispersal syndromes of plant species can be limiting, as these relationships can be habitat specific. Although we used the best data available from a trait database and field guides, we recognise that next steps should include an improvement on data used. These data include plant-insect associations for additional insect taxa (e.g., wild bees as pollinators or plant-and leafhoppers as herbivores), and of improved occurrence data (for our work, no data on the altitudinal occurrence of moths was available). Local entomological specialists can help to compile realistic lists of plant-insect interactions, and we postulate that this method could also make excellent applied use of pollinator networks or food web data across trophic levels.

Here, targets were set to ensure a minimum of one attribute was present in solutions, to allow for direct comparison between objectives, but this means other species were categorized as redundant when not selected. In practice, including an abundance of targeted attributes within solutions is desirable and likely beneficial for restoration outcomes, and these targets can be adapted for various needs.

Similarly, we assume that the cost of including a plant species is the same to test our questions independent of costs, but the approach can and should account for cost variation to acquire, store and reproduce seeds as this will likely hold great influence on prioritized solutions in practice (Jiménez-Alfaro et al., 2020). Reporting on the costs of conservation and management actions is largely inadequate and non-standardized (Iacona et al., 2018), and we know from previous research that only a small proportion of seeds are usually available for purchase (Ladouceur et al., 2018). Prioritization approaches could also guide future efforts for seed supply and policy by informing collection, farming and storage for an expanded restoration species pool. Further, including these real costs in decision-making frameworks can help to plan efficient projects.

### Ways forward and Conclusions

This proof of concept is the first step towards framing future empirical research in ecological restoration of natural ecosystems. We call for empirical field tests for this approach to take place, which will require bringing together interdisciplinary collaboration across subfields of ecology and conservation. We provide a transparent and robust approach that could move restoration efforts towards prioritizing plant species to maximise targets and minimise costs offering quantitative decision-making support. This approach could be applied to any system and/or targets which could also contribute to many stages of restoration decision-making and could play an important role in delivering efficient, targeted solutions. However, similar approaches will need robust ecological data to be applied to specific cases studies and restoration targets, preferably at regional or local scales.

## Acknowledgements

E.L., C.B. & B.J.A. acknowledges the support of the research leading to these results has received funding from the People Programme (Marie Curie Actions) of the European Union’s Seventh Framework Programme FP7/2007–2013 under REA grant agreement no. 607785, as a part of the NAtive Seed Science TEchnology and Conservation (NASSTEC) Initial Training Network (ITN). E.L., R.vK. & B.J.A. gratefully acknowledge the support of iDiv funded by the German Research Foundation (DFG–FZT 118, 202548816). E.L. is grateful for the support of the Alexander von Humboldt Foundation.

## Data Availability

All data used for these analyses will be made available open access at FigShare and Code to reproduce analyses will be made available on GitHub/Zenodo.

## Author Contributions

P.H., H.P. and E.L. conceived the idea. E.L. and B.J.A designed the case study, defined the species pool, and made a data collection plan. E.L., D.S., and R.vK. collected and requested data. P.P. and J.H.C. donated plant trait data. E.L. performed the analysis. J.M., H.P. and P.H. provided guidance on analyses and interpretation. E.L. and J.M. wrote the manuscript. All authors contributed to editing and shaping the manuscript into the final version.

## Supplementary Information

## Appendix 1: Special Indications and Expanded Methods

### Non-Spatial Prioritizr or MARXAN use

The framework we developed draws from the ‘minimum set’ problem definition commonly applied to spatial conservation planning activities which rely on decision-support tools that provide algorithmic solvers, like MARXAN or prioritizr. In this context, spatially-defined areas of the land or water are the planning units. The algorithms aim to meet target amounts for important attributes found in the planning units. These attributes are commonly referred to as conservation features (Game & Grantham 2008; Hanson et al. 2019). In the non-spatial use of these tools, demonstrated here for ecological restoration, the plant species become the planning units and the attributes become the conservation features. This requires nothing more than arranging the data in the appropriate manner. Instead of a site by feature matrix, the data becomes a plant species by attribute matrix.

Using these tools requires the development of four types of input variables in the non-spatial context for the selection of plant species for ecological restoration planning. First, the identification of a suite of desirable species that might be encompassed in the entire species pool, potentially all desirably occurring in the target habitat of choice. We give more description of this step below under ‘Determining Species Pools’. Second, a plant attribute value for each plant species is the numerical or categorical attribute value associated with each plant under every single attribute field, which can be represented as present (1) or absent (0). Each attribute must have its own column. For example, when using an attribute such as ‘flowering duration’, every month of the year must be represented as a single attribute, and a plant species can have a presence value in the months where it flowers. If a plant species has a relationship with a specific species of pollinator in terms of both a nectar resource, and a larval diet resource, there would be an attribute column represented for each pollinator species, and for each type of ecological relationship represented. See Table S1 for how this is represented in the species-attribute data matrix practically.

Third: targets are identified and set that specify the quantities of each attribute that should be represented in the final selections. These targets serve as an initial hypothesis for testing necessary levels of replication and abundance to ensure attribute presentation (Chan et al. 2006). Targets can be set for different representation levels, or specific ‘units’ can be eliminated from solutions, or forced into solutions based on different controls. Lastly, costs are set, where a numerical value can be assigned to each planning unit. Each species was given a value of 1 for this proof-of-concept exercise, but further work can be done incorporating the real cost of hand collecting, buying seed or plugs for plantings and these costs could be compared in the prioritizations. This would add further complexity to the decision support process, but increase the quantitative power and value of this exercise.

Once all variables are set, software use instructions would be followed (Game & Grantham 2008; Ball & Possingham 2009; Hanson et al. 2019), and MARXAN or prioritizr is run to select priority plant species that collectively constitute a selection that captures all desired target attributes and function within the minimum amount of plant species. For every objective, we ran prioritizr 100 times producing one hundred solutions for every objective-the standard best practice amount. These selections act as replicates for quantitative support in decision making, and ensure comparability between objectives. prioritizr identifies a single optimal solution which is the plant species selection that meets targets and minimises cost, but can be run with a portfolio option to generate more than one solution. Because we gave all plant species equal ‘cost’ values in this exercise as a proof of concept, the problems had simpler solutions than if the plant species were given real costs associated with the purchase or hand collection of wild seed. As a result, many solutions contained different constellations of plant species, but received equally high scores. We always used the first solution as identified by prioritizr for analyses. However, in Figure 4, where we refer to the ‘solutions with the best score’ this refers to the fact that we used all solutions with equally high scores for this analysis. Users of this method may encounter similar conditions when running problems where all plant species have been given an equal cost, depending on the species richness of the entire species pool considered, the targets, the nature of the attributes, and the constraints of a project.

### Determining Species Pools

Methods for determining the appropriate target habitat type for restoration, and species pool for restoration species mixes vary by region, and when applying this method to different areas, these standards can be used to determine this appropriate species list. If a practitioner were starting with a restoration site that already had some species present, or had natural regeneration potential (eg. from the seed bank, or successional species able to natural recolonize) the site should be surveyed and assessed for this first, local expert knowledge used, and these species can still be included in species lists, but eliminated or constrained within solutions (see MARXAN documentation). Clearly identifying the set of species appropriate to be used for restoration species lists has a major impact on this process and resulting solutions and this step should be approached carefully and appropriately to each project.

**Figure S1:**
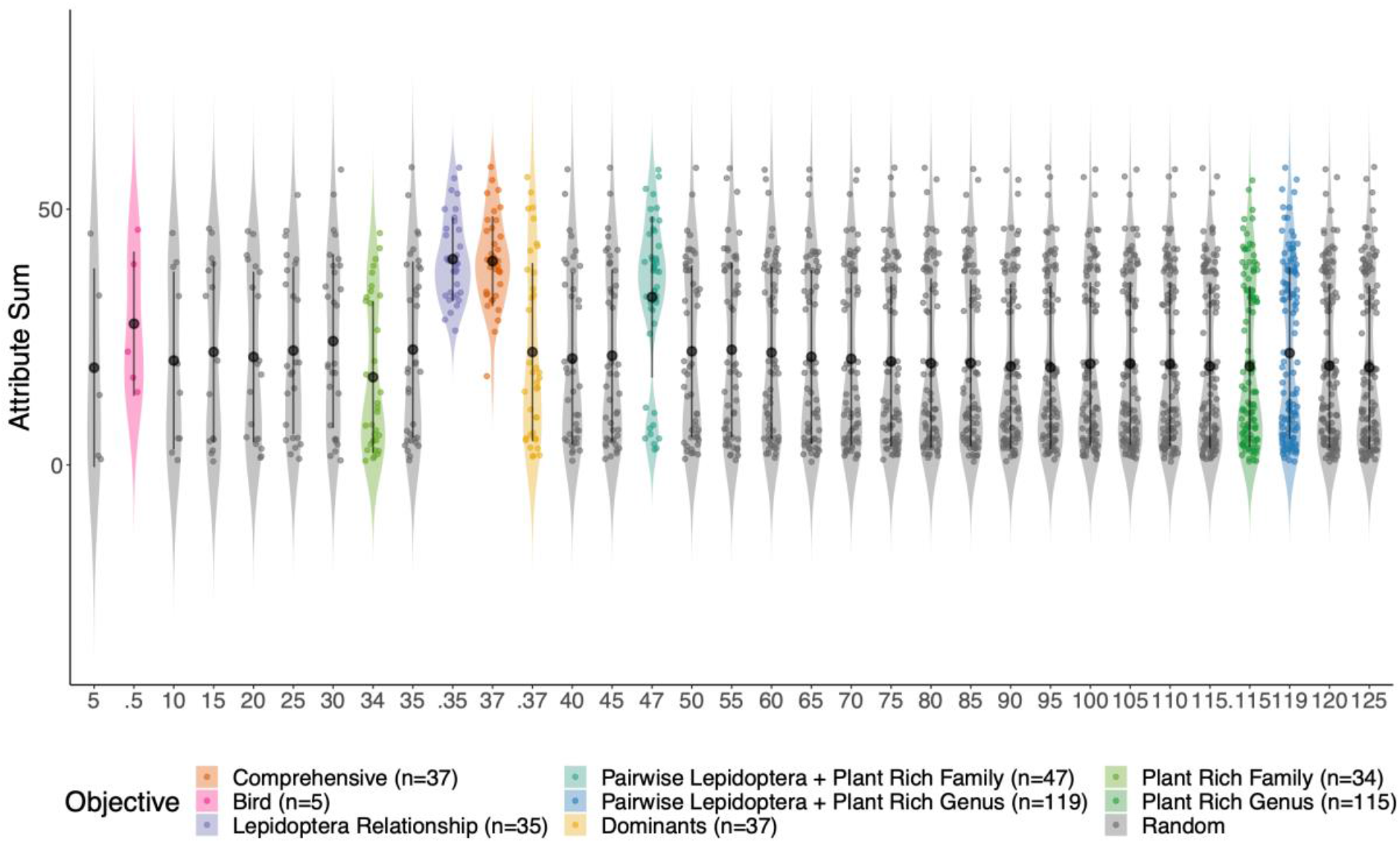
This is an expanded version of Figure 2. The attribute sum (total number of attributes per plant species) captured by each plant species in the best solution of every objective. The number of species in each solution is labelled along the x-axis. Colors represent each objective, and black is the species chosen at random for every step of number of species. Each point is a plant species. The black points represent the mean and the bars represent the standard error around the mean. The shaded area represents the spread and the density of the data. n: number of plant species in the best solution of every objective.

**Figure S2:**
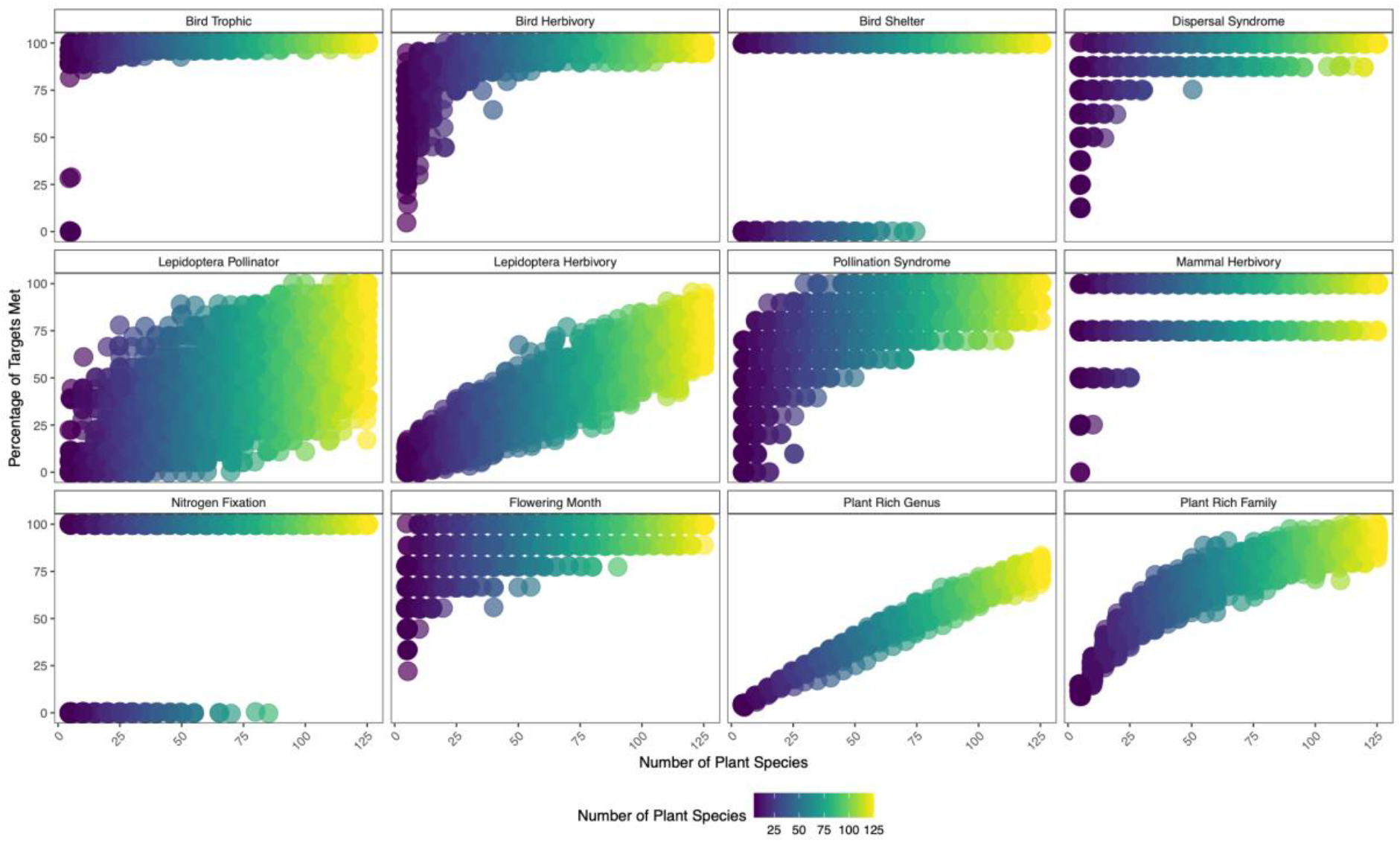
A Trade-off plot showing the trade-off between the number of plant species randomly selected, and the corresponding proportion of targets (representing 1 of each attribute within a category) for each broad group of ecosystem function or service provided. Plant species were selected randomly 100 times for each level in intervals of 5 from 5 plants species increasing to 125 plant species.

